# Functional profiling of the male gametophyte-specific promoter fragment of *Arabidopsis PIRL6* gene and prediction of *cis*-regulatory elements

**DOI:** 10.1101/2024.01.09.574950

**Authors:** T.G. Ajith, Jasmine M. Shah

## Abstract

Gametophyte-specific promoters drive the expression of genes in male and/or female gametophytes. These have applications in breeding experiments, gene function identification, developmental biology-related studies, and off-lately in genome editing also. The *Plant Intracellular Ras-group Leucine-Rich-Repeat* 6 (*PIRL6*) gene is known to be necessary for male and female gametogenesis. Using the *GUS-* based deletion analysis, we have identified the *PIRL6* promoter length that is essential for exclusive expression in the male gametophyte of *Arabidopsis thaliana.* We studied the strength of various lengths of *PIRL6* promoters in different tissues, by GUS expression quantification. The male-gametophyte-specific promoter segment (PA1) exhibited stronger expression in mature anthers than the younger ones. We identified 50 other genes that co-expressed with *PIRL6* in *Arabidopsis* using the Expression Angler tool. Gene ontology (GO) analysis shows that these 51 co-expressing genes were predominantly involved in cell differentiation. By comparing the promoter sequences of these 51 genes, the presence of three over-represented known motifs, POLLEN1LELAT52, ACGTATERD1 and CIACADIANLELHC was identified. We could also predict the presence of three novel *cis-*regulatory elements in the co-expressing gene network using the MEME suite tool. Additionally, we confirmed the functionality of the ABRE and P-box elements present in *PIRL6* promoter using the tobacco leaf transient assay. Thus, we cloned and functionally confirmed the promoter region of *PIRL6* required for male gametophyte-specific expression, compared this promoter with those of 50 other co-expressed genes and predicted their functions, and analyzed their *cis* regulatory regions and predicted three novel motifs also.

**Key message:** A pollen-specific promoter fragment of *PIRL6* was functionally characterized using *GUS*-based deletion analysis. Fifty other co-expressed genes were compared and novel *cis*-regulatory elements were predicted. Two hormone-responsive elements in the *PIRL6* promoter were found to influence the *GUS* expression. PA1 promoter can be used in experiments that require male gametophyte-specific expression of genes.

## Introduction

The life cycle of fertile angiosperms generally have two stages of ploidy, the gametophytic (haploid) and sporophytic (diploid) (Goldberg et al. 1993). In contrast to lower plant species where the gametophyte is the dominant and a free-living generation (Cove and Knight, 1993), gametophytes of angiosperms are smaller and less complex than the sporophyte. The female and male gametophytes of angiosperms are formed within specialized organs of the flower, the ovary and anther, respectively. While these gametophytic life cycles begin with the meiotic division of their respective mega or microspore mother cells in the ovary and anther, respectively, the sporophytic stage begins with the fertilization of gametes produced from the gametophytes. As a consequence of meiosis, the sporophyte produces two types of spores, the megaspores and microspores, from which the female (embryo sac) and male (pollen) gametophytes develop and produces the sex cells or, gametes or, egg and sperm, respectively, which fuse during fertilization and restore the ploidy. Gametophyte development from the sporogenous cells requires a coordinated expression of many genes in both, the haploid gametophytic cells, and the sporophytic tissues. Expression studies at the whole-transcriptome level have been done in both male (Deveshwar et al. 2011; Rutley and Twell 2015; Twell 1998) and female (Drews et al. 2010; Rutley and Twell 2015; Yu et al. 2005) gametophytes and, even in sperms (Borges et al. 2008), of various plants. Thousands of genes that exhibit differential expression in angiosperm gametophytes have been identified. There are a few genes such as *AtPTEN1* of *Arabidopsis* (Gupta et al. 2002), *Pex1* of *Zea mays* (Rubinstein et al. 1995), and *Bnm1* of *Brassica napus* (Treacy 1998), which are reported to express only in the haploid generation of plants, the gametophyte only, and not the sporophyte. Such genes behave differently in gametes because of their regulation, which include the special category of tissue-specific promoters known as gametophyte-specific promoters.

Tissue-specific promoters are known to control the expression of genes spatially and temporally in all organisms. Hence, when such promoters are used to regulate the expression of the gene of interest *in trans*, they can avoid potential negative effects of a constitutive promoter such as metabolic burden (Ye 2000). Tissue-specific promoters have key importance in the field of plant metabolic engineering (Ye 2000). Such promoters have high demand in facilitating the compartmentalized production of desirable proteins of pharmaceutical importance from organs such as seeds (Kusnadi et al. 1998), leaves (Komarnytsky et al. 2000), roots (Borisjuk et al. 1999), or fruits (Daniell et al. 2001). He et al. (2018) reported the use of embryo-specific promoters to drive a gene in order to generate transgene-free genome edited rice plants. Studying the behavior of promoters with the aid of reporters such as GFP and GUS has also added to a better understanding of the functions of the respective genes under their control (De Ruijter et al. 2003). Gametophyte-specific promoters, which target the expression of genes in male/female gametophytes are deployed in breeding experiments, gene function identification, and developmental biology-related studies (Oo et al. 2014, Dwivedi et al. 2010). Wang et al. (2015) used female gamete-specific promoter to enhance the generation of homozygous genome edited *Arabidopsis* plants. Gametophyte-specific promoters can be categorized as those which express in both, the sporophytic tissue and the gametophyte, and, those which express only in the gametophyte or the haploid generation of plants (Drouaud et al. 2000).

There are previous reports of functional characterization of promoters specific for the male gametophytic generation. These include the promoters for the genes *LAT52* and *LAT59* from tomato (Twell et al. 1991), *OSIPA* (Swapna et al. 2011) and *OSIPK* (Gupta et al. 2007) from rice, *AtVEX,* and *AtGEX2* (Engel et al. 2005), *HAP2* (von Besser et al. 2006), *DUO3* (Brownfield et al. 2009), and *WRKY34* (Zou et al. 2010) from *Arabidopsis* and many others as reviewed in Smirnova and Kochetov (2020). The reports on female gametophyte-specific promoters are very few in comparison to their male counterparts. Some of these include those driving the genes *At5g40260* of *Arabidopsis* and *BRA0029160* of Chinese cabbage (Kim et al. 2013), and *EC1.2* of *Arabidopsis* (Wang et al. 2015, Steffen et al. 2007). Based on the expression of tissue-specific genes, more promoters are being functionally characterized. Alongside, novel organ-specific motifs, conserved motifs, and other regulatory elements in such promoters are also being discovered (Hernandez-Garcia and Finer 2014; Smirnova et al. 2019).

The *Plant Intracellular Ras-group Leucine-Rich-Repeat* (*PIRL*) is a family of genes exhibiting gametophytic expression (Forsthoefel et al. 2013). These code for leucine-rich repeat proteins (LRRs) capable of initiating particular protein-protein interactions to facilitate signal transduction pathways. The *PIRL*-encoded LRRS are structurally related to Ras-group LRRs, which are the LRRs involved in Ras (rat sarcoma) signal transduction pathways in animals, and hence PIRLs are grouped under the Ras-group LRRs (Forsthoefel et al. 2005). One member of this family of genes, the *PIRL6*, was found to be necessary for both male and female gametogenesis and alternative splicing negatively regulates its sporophytic expression (Forsthoefel et al. 2018). In this report, we have studied the functions of various deletion fragments of the *PIRL6* promoter and identified the region essential for pollen-specific expression. We have predicted predominant *cis*-elements in the promoters of *PIRL6* and the genes that co-expressed with it. We have also confirmed the function of two hormone-responsive motifs in the *PIRL6* promoter.

## Materials and methods

### DNA extraction, primer designing, PCR, and sequencing

The total DNA of the *A. thaliana* Columbia ecotype was extracted using a modified CTAB-based method (Rogers and Bendich 1998). All primers were designed using the tool Primer Quest and purchased from G-Biosciences. All PCR reactions were performed using the PCR components from Takara bio Inc. The annealing temperature of each reaction varied according to the respective primer pair (Table 1).

### Restriction digestion, ligation, bacterial transformation

The restriction enzymes *EcoR*I and *Pst*I, and the ligation mix, were purchased from New England Biolabs. Both digestion and ligation setups were as per their respective manufacturer’s instructions. *E. coli* strain DH5α was transformed with the desirable clones using heat shock method (Froger and Hall 2007). The transformation of *Agrobacterium tumefaciens* strain GV3101 with these clones was by freeze-thaw method (Weigel and Glazebrook 2006).

### *Arabidopsis* transformation

*Arabidopsis thaliana* plants from the Columbia ecotype were transformed with the *Agrobacterium tumefaciens* strain GV3101 harboring the desirable clone, using the floral dip method (Clough and Bent 1998). The putatively transformed seeds were selected on MS media plates containing 30 μg/ml hygromycin.

### GUS histochemical staining and fluorometric assay

GUS histochemical staining of plants was as per the protocol of Jefferson (Jefferson et al. 1987). Subsequent chlorophyll removal of the tissues was performed by overnight treatment with 70% ethanol. These tissues were observed under the stereo microscope (Carl Zeiss Suzhou) and photographed.

GUS fluorometric assay was performed as per the protocol of (Jefferson et al. 1987). The total protein concentration in the homogenate was calculated using Bradford’s method (Bradford 1976). The GUS fluorometric activity was assessed based on the nmol of 4-MU produced per minute per milligram of protein. One-way analysis of variance (ANOVA) was performed to determine statistical significance for relative gene expression among different transgenic and non-transgenic plant samples. Means were compared using Tukey honestly significant difference (HSD) post hoc comparisons.

### Promoter sequence analysis

The complete sequence of the *PIRL6* promoter was analyzed and the distribution of motifs was studied using the online tool PlantCare.

### Coexpression analysis

The co-expression network of *PIRL6* was generated with the Expression Angler tool (Toufighi et al. 2005). The top 50 coexpressed genes in the ‘Custom Expression Pattern’ as defined by *PIRL6* in the ‘Microgametogenesis - Tissue Specific view’ were downloaded.

### Pathway visualization analysis

Using Expression Angler’s top 50 coexpressed genes with the same ‘tissue-specific‘ expression pattern as *PIRL6*, enriched Gene Ontology (GO) terms were discovered [reference genome: Affymetrix ATH1 Genome Array (blast); test: hypergeometric test with Hochberg FDR correction] (Du et al. 2010). A cutoff of mapping to at least three terms was chosen since it produced results that were more informative (instead of the default of five). Evidence for the GO annotations enriched in this set was examined by entering the coexpressed gene list into the g:Profiler programme (Reimand et al. 2016). Statistical domain scope was set to ‘Custom over all known genes’ backdrop set to ’AFFY ATH1 121501’, and significance threshold set to ‘Benjamini-Hochberg FDR’ as closely as possible to the AgriGO settings. *A. thaliana* was chosen as the species, and all other variables were kept at their default values.

### Cistome analysis and *de novo* motif prediction of *cis*-regulatory element

The Cistome tool was used to look for common motifs in the 51 ‘tissue-specific’ coexpressed genes from the Expression Angler tool (Austin et al. 2016). The TAIR Upstream (TrSS) 1000 bp promoter data collection was used, and for Step 3 ’Paste in your own PSSMs or consensus sequences and/or choose precomputed motifs from other sources for *de novo* mapping was used, with ‘All PLACE elements’. Ze cutoff > 2 Significant Motifs were the only ones chosen to be displayed (instead of the default 3, other defaults were kept as such).

One kilobase upstream sequence flanking the ATG start codon was collected from 51 co-expressed genes. We subjected these promoter sequences to the command line application of ‘Multiple Expectation maximizations for Motif Elicitation’ (MEME) Suite (v5.0.5) for *de novo cis*-element prediction, overrepresented in the gene network. The algorithm was used to look for motifs with lengths of 6 to 50 nucleotides with E values less than 1. The ‘zero or one occurrence per sequence’ (zoops) option was used to conduct motif searching for each set of upstream regions.

### Agro-infiltration of tobacco leaves and hormone treatment

Four-week-old *Nicotiana tobacco* leaves were infiltrated with the appropriate *Agrobacterium* culture or plain infiltration media (Ali et al. 2018). The overnight-grown *Agrobacterium* culture was resuspended in induction media (Mangano et al. 2014) and allowed to grow at 28°C for 6 hours in a shaker. The bacterial pellet was collected and resuspended in infiltration media (10 mM MgSO4, 10 mM MES). The infiltration was done on the abaxial side leaves using 1 ml syringes without needles. The entire plant was then covered with plastic bags and kept in the growth chamber at 22°C with 16 and 8 hours of light and darkness, respectively. The next day, the plastic bags were removed and the plants were left in the same conditions for two more days. After 48 hours of agro-infiltration, 200 μM each of ABA or GA3 (Sigma Aldrich) were sprayed independently on the leaves. After 24 hrs, the leaf discs comprising the infiltrated area were collected and subjected to GUS histochemical staining and fluorometric assay.

## Results

### Cloning variable lengths of gene promoter

To functionally characterize various promoter regions upstream of the *PIRL6* gene, three different lengths of promoter sequences, 1128 bp (-1111 to +17), 762 bp (-745 to +17), and 646 bp (-629 to +17), also known as PA1, PA2, and PA3, respectively (Fig. 1) were PCR amplified using different primer pairs (S. Table 1). The 5’ end of all three forward primers (F1, F2, and F3) were designed to carry an *EcoR*I site. Similarly, the 3’ end of their common reverse primer (R) had the *Pst*I, site. Thus, the PCR-amplified products were double-digested using these two restriction enzymes, and cloned into the *EcoR*I*/Pst*I sites located upstream of the promoter-less GUS gene of the binary vector pCAMBIA 1381 (S. Fig.1). The *GUS* gene is a reporter that encodes the enzyme β-glucuronidase. Three putative clones were generated, namely pAJPIRL1, pAJPIRL2, and pAJPIRL3 harboring PA1, PA2, and PA3 respectively, after transformation into *E. coli*. After confirmation by restriction digestion and sequencing, all three clones were independently transformed to the *Agrobacterium* strain GV3101.

**Fig. 1.**
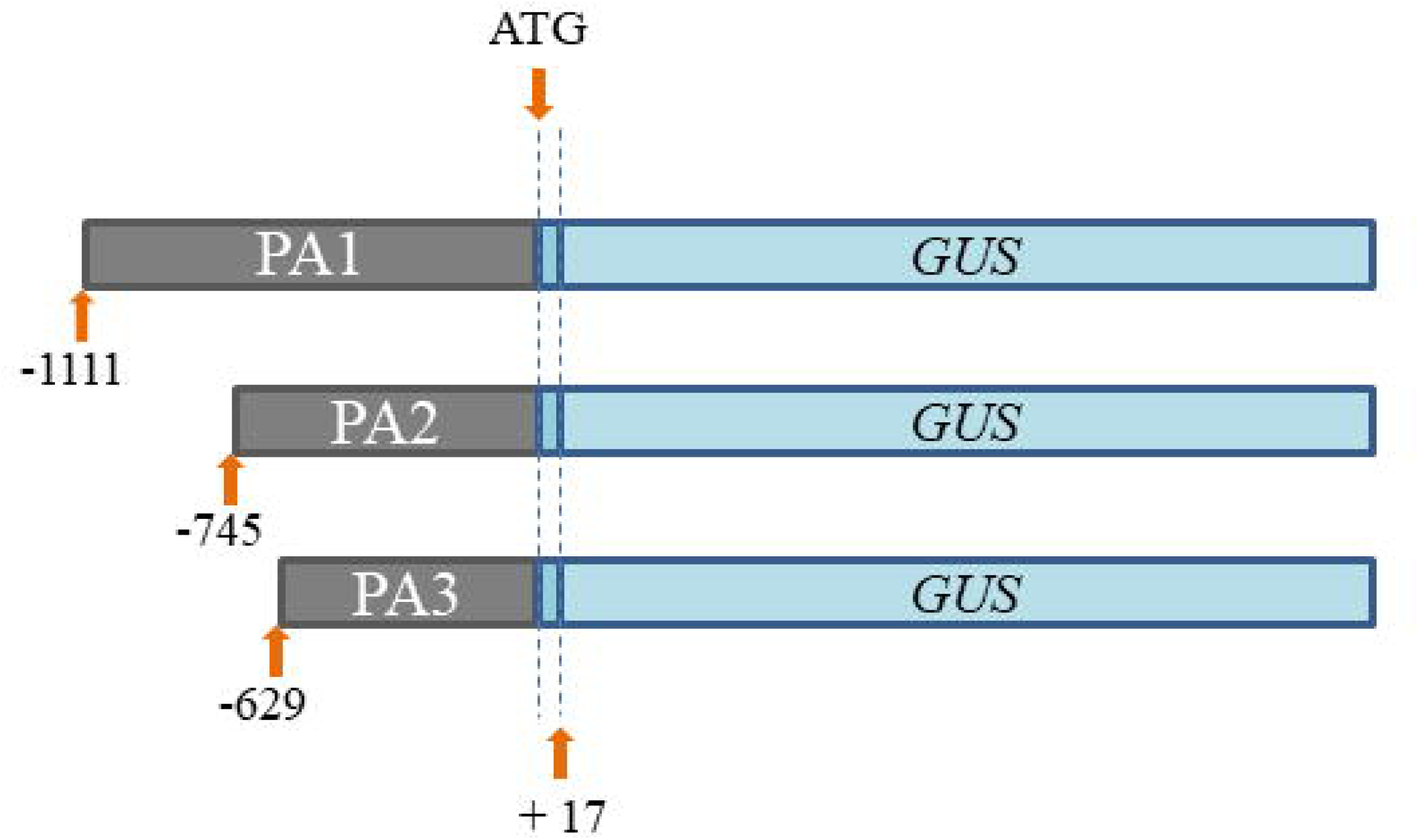
Deletion analysis of *PIRL6* promoter. PA1, PA2 and PA3 are three different lengths of *PIRL6* promoter fused to *GUS* gene. The negative number upstream region of the promoter indicates the starting site of the promoter. The downstream end was at +17 for all the three promoters.

### *Arabidopsis* transformation and deletion analysis of *PIRL6* promoter

For each of the three deletion constructs, at least 15 hygromycin-resistant plants were obtained. To begin with, five putatively transformed T0 plants from each transformation were confirmed using PCR with *GUS* primers (S. Fig. 2); if required, more were screened. Four single-copy events from each transformation were identified by subjecting the T1 seeds to segregation analysis on hygromycin-containing media. The T2 plants from each line were selfed and their zygosity was analyzed by repeating the hygromycin-sensitivity test. T3 seedlings were collected only from homozygous T2 plants that generated all hygromycin-resistant progeny. In order to see the influence of various promoter lengths on the expression of the *GUS* gene in various organs, T3 plants were subjected to GUS staining at various stages such as seedling, vegetative, as well as flowering. While the GUS staining was seen only in anthers of the plants derived from line PA1 (Fig. 2a), staining of anthers as well as other sporophytic tissues, such as leaves, stems, and sepals were seen in plants derived from lines PA2 and PA3 (Fig. 2b). Petals and ovaries from plants derived from all three lines PA1, PA2, and PA3 did not show GUS staining. As the anthers comprise of two components, the sporophytic (anther wall) and gametophytic (pollen grains), the pollen grains were separated from the anthers and examined (S. Fig. 3). This revealed that the anther walls did not stain blue, and the anthers appeared blue because of the stained pollen grains, thereby indicating that the PA1 promoter was pollen-specific. Of the three promoters, PA1 was the longest (1128 bp), followed by PA2 and PA3 (762 bp, and 646 bp, respectively).

**Fig. 2.**
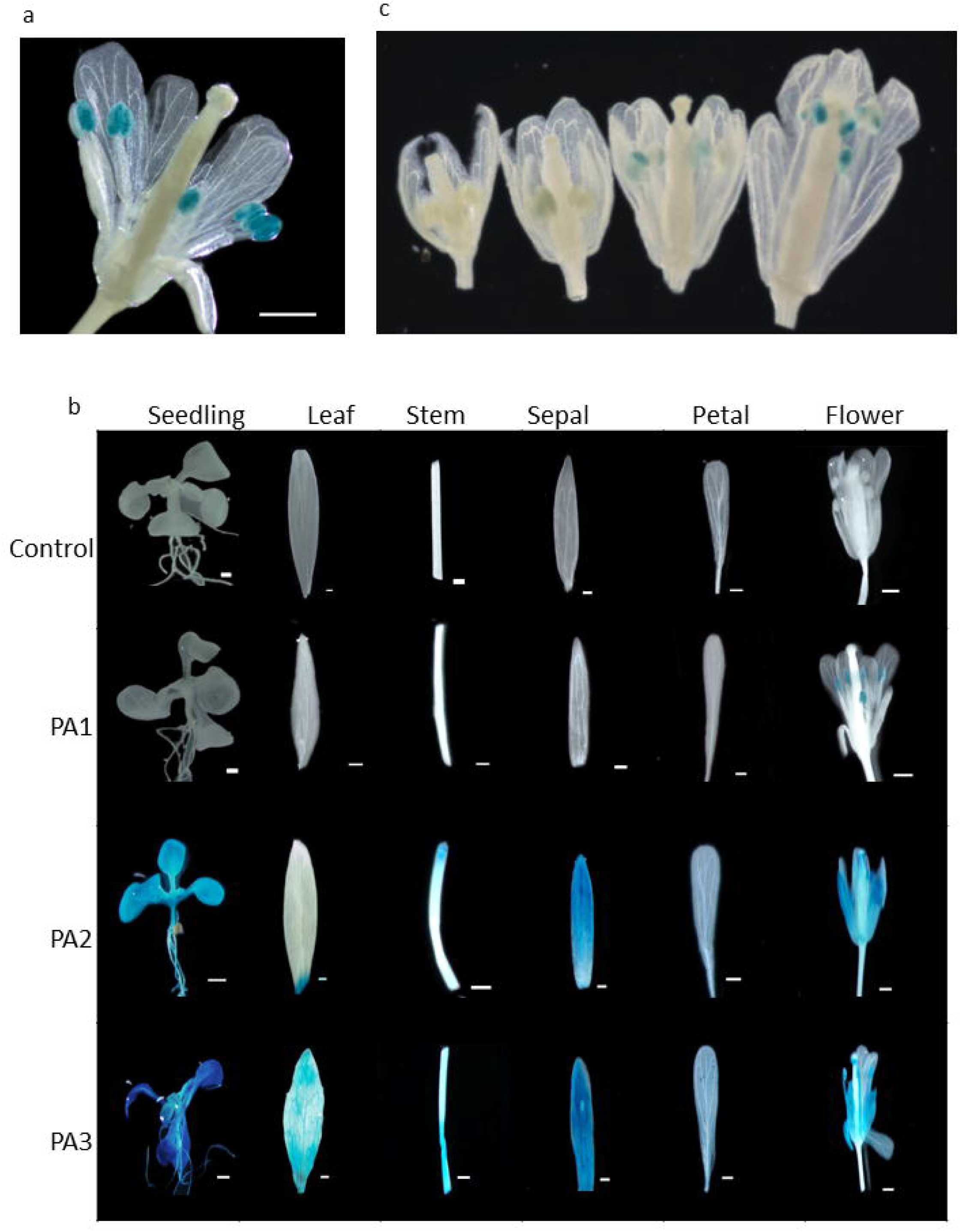
GUS expression pattern in *Arabidopsis* plants transformed with different lengths of *PIRL6* gene promoter. (a) Flower of plant transformed with *GUS* under PA1 promoter, showing GUS staining in the anthers. (b) GUS staining of various tissues of wild (control) and transgenic *Arabidopsis* plants harboring different promoter::*GUS* cassettes of PA1, PA2 and PA3. Bars = 1mm. (c) Evaluation of GUS activity under the control of PA1 promoter at different stages of *Arabidopsis* flower development, from left to right – un opened flower bud, one day after the opening of flower, three day after opening of flower, five day after opening of flower.

In order to study the temporal behavior of PA1 promoter, flower buds of different stages, and opened flowers were subjected to GUS staining. While the younger buds did not show any blue coloration, anthers from buds one day before opening showed mild blue staining, and those in completely opened flowers stained darker (Fig. 2c). This revealed that the GUS expression was predominantly in the mature pollen grains compared to that of developing pollen grains.

### Quantification of GUS expression

To gauge the intensity of GUS expression in various tissues due to different promoter lengths, GUS fluorometric analyses were performed. Various organs such as leaf, stem, sepal, petal, gynoecium, mature anther, and young anther, from five different homozygous plants derived from each of the lines PA1, PA2, and PA3 were analyzed and all these showed varied levels of GUS expression (Fig. 3). Unlike the PA1 plants, which had higher expression in anthers (Fig. 3a and b), the PA2 and PA3 plants exhibited significantly higher GUS expression in leaves and other sporophytic tissues (Fig. 3). In comparison to the untransformed plant, all organs from the plants derived from all three transformed lines exhibited faint expression, which was not visible microscopically. Nevertheless, the average GUS activity from the anthers (young and mature) was six-fold higher than that from other sporophytic tissues (leaf, stem, petal, and sepal) of PA1-derived plants.. In contrast to PA1, the average GUS activity of anthers from PA2- and PA3-derived plants was 6.3 and 4.2 fold lesser than the sporophytic tissues (leaf, stem, and sepal). Thus, it was confirmed that for *PIRL6* to express in male gametophytes, the PA1 promoter region was essential.

**Fig. 3.**
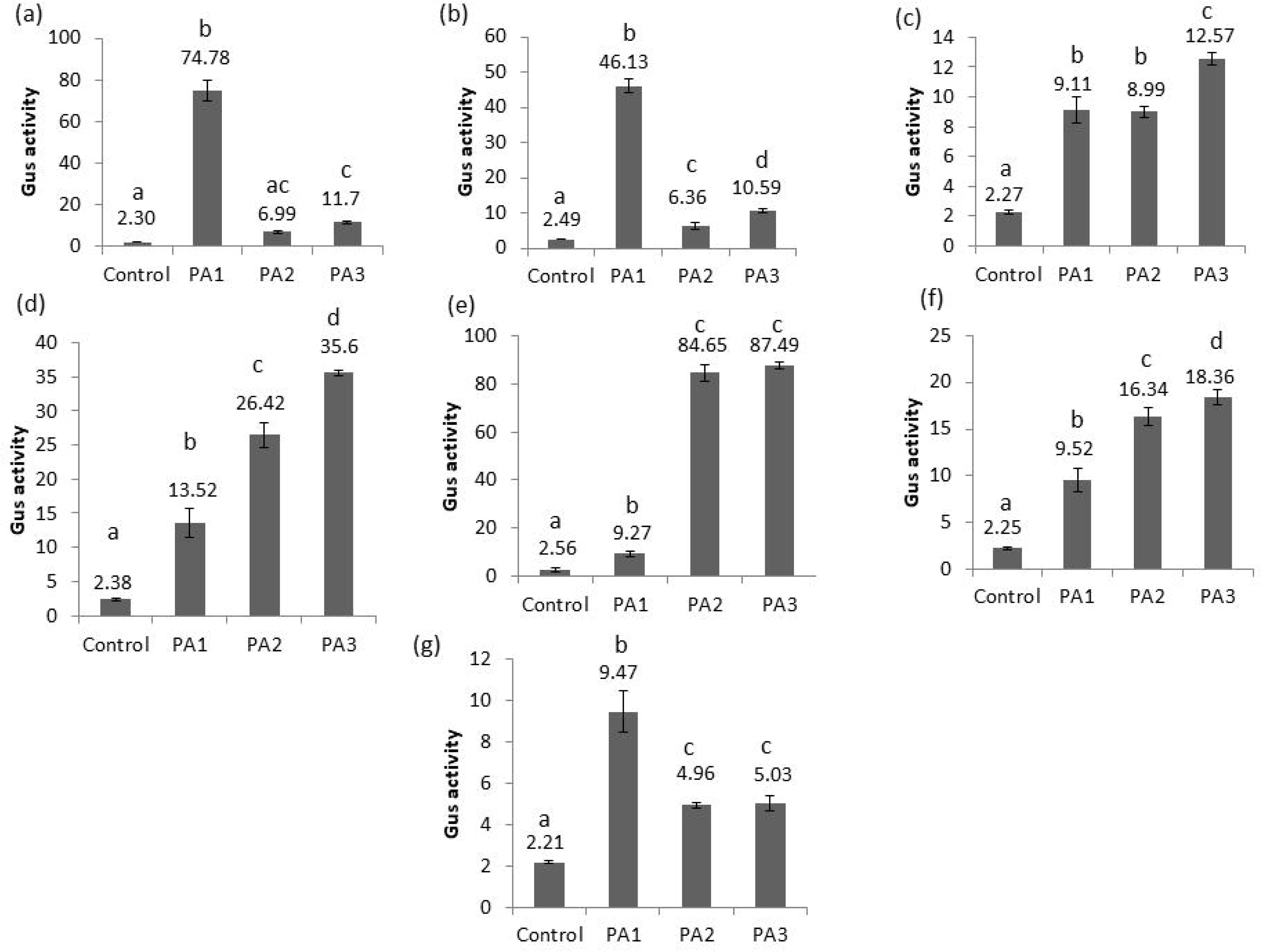
Quantitative measurement of GUS activity in different tissues of transgenic *Arabidopsis* plants harboring PA1, PA2, PA3 promoter cassette and wild plant (control). (a) Mature anther, (b) Young anther, (c) Gynoecium, (d) Stem, (e) Leaf, (f) Sepal, (g) Petal. The error bars indicate ± Standard deviation of the means of four biological replicates. The symbols a, b, c and d denote a significant difference between the respective groups at p < 0.01, if not represented by the same symbol. GUS activity was assessed based on the nmol of 4-MU produced per minute per milligram of protein.

### Functional prediction of the co-expressed genes of *PIRL6*

It could be possible that the region exclusive for PA1 harbored male gametophyte-specific regulatory sequences. In order to get more information about such sequences, it is required to study promoters of other genes that co-expressed with *PIRL6*. Towards this end, top 50 genes that co-expressed with *PIRL6* in male gametophyte were predicted (S. Table 2) using the Expression Angler tool. To determine the likely function of the *PIRL6* co-expressed gene network, GO analysis was performed using two different functional classification tools, the Agrigo and g:Profiler. While Agrigo is good at prediction of various possible functions, g:Profiler displays GO functions with the source of experimental evidence. Functions predicted by ‘AgriGo’ include cell differentiation, regulation of anatomical structure size, regulation of biological quality, and regulation of cellular component size (Fig. 4). The g:Profiler analysis resulted in the prediction of two possible functions, the regulation of pollen tube growth, and regulation of cell morphogenesis involved in differentiation, based on the previously known evidence from three co-expressed genes, AT4G13240, AT1G77980, and AT1G18750. Nevertheless, cellular differentiation was the commonly predicted function by both the tools for GO analysis.

**Fig. 4.**
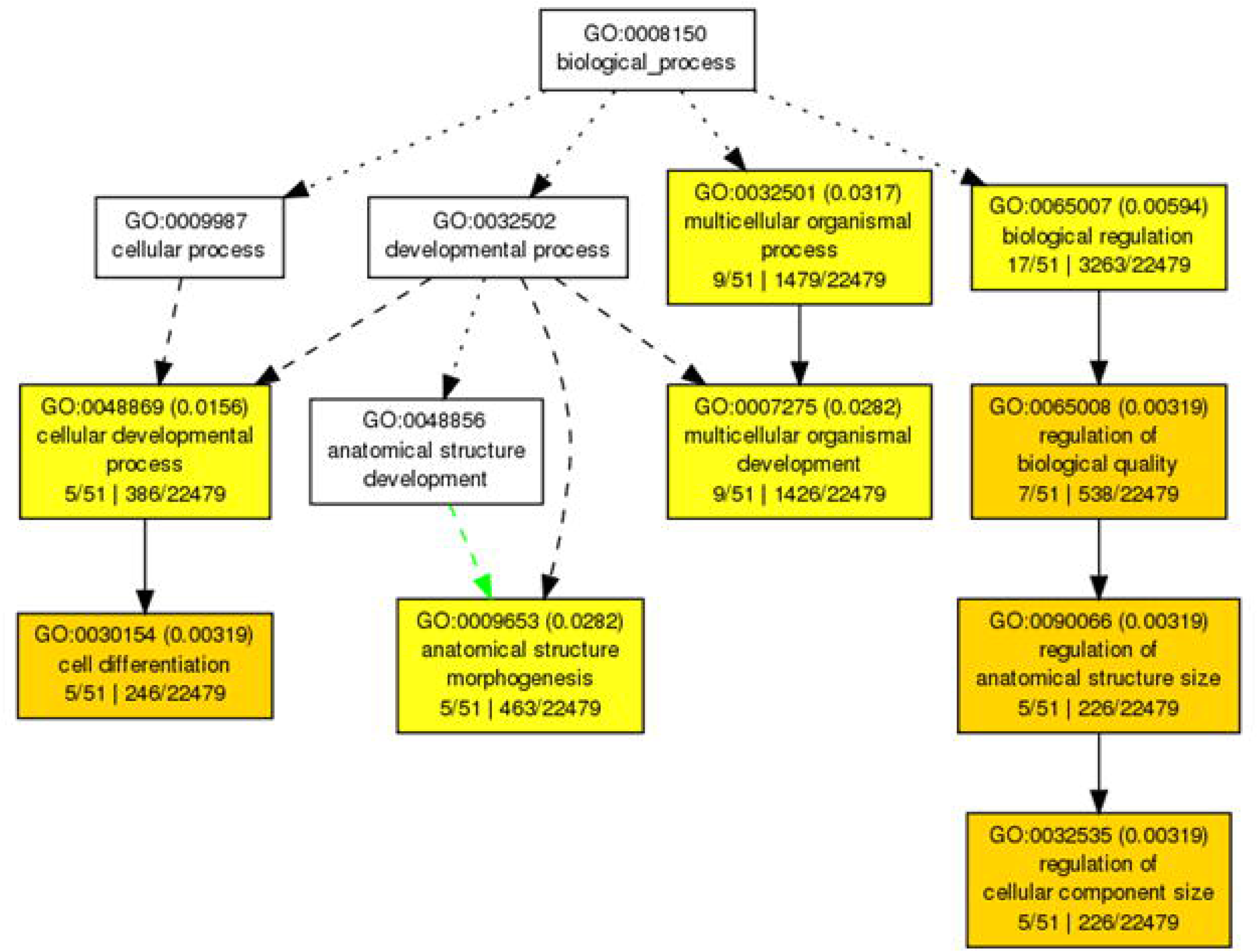
AgriGo Gene Ontology (GO) analysis of the 50 coexpressed genes of *PIRL6* significantly enriched for the GO BP “cell differentiation”. The graph has boxes that represent GO terms and are labelled with their GO IDs, term definitions, and statistical data. Significant terms are shown by coloured boxes, whereas non-significant terms are displayed as white boxes (adjusted P 0.05). The diagram shows a positive correlation between the term’s level of enrichment and the colour saturation of a box. Solid, dashed, and dotted lines represent two, one and zero enriched terms at both ends connected by the line, respectively. The graph is set to have a top-to-bottom rank order.

### Promoters of co-expressed genes are enriched with the POLLEN1LELAT52 motif

Cistome analysis (S. Fig. 4) of the promoters of *PIRL6* and all the above 50 co-expressed genes showed the predominant presence of three known motifs, POLLEN1LELAT52, ACGTATERD1 and CIACADIANLELHC (Fig. 5a). A total of 628 motifs were generated, out of which 438, 137 and 53 belonged to POLLEN1LELAT52, ACGTATERD1, and CIACADIANLELHC, respectively. POLLEN1LELAT52 was in variably present, as well as was most predominant, in all the genes analyzed (S. Fig. 5).

**Fig. 5.**
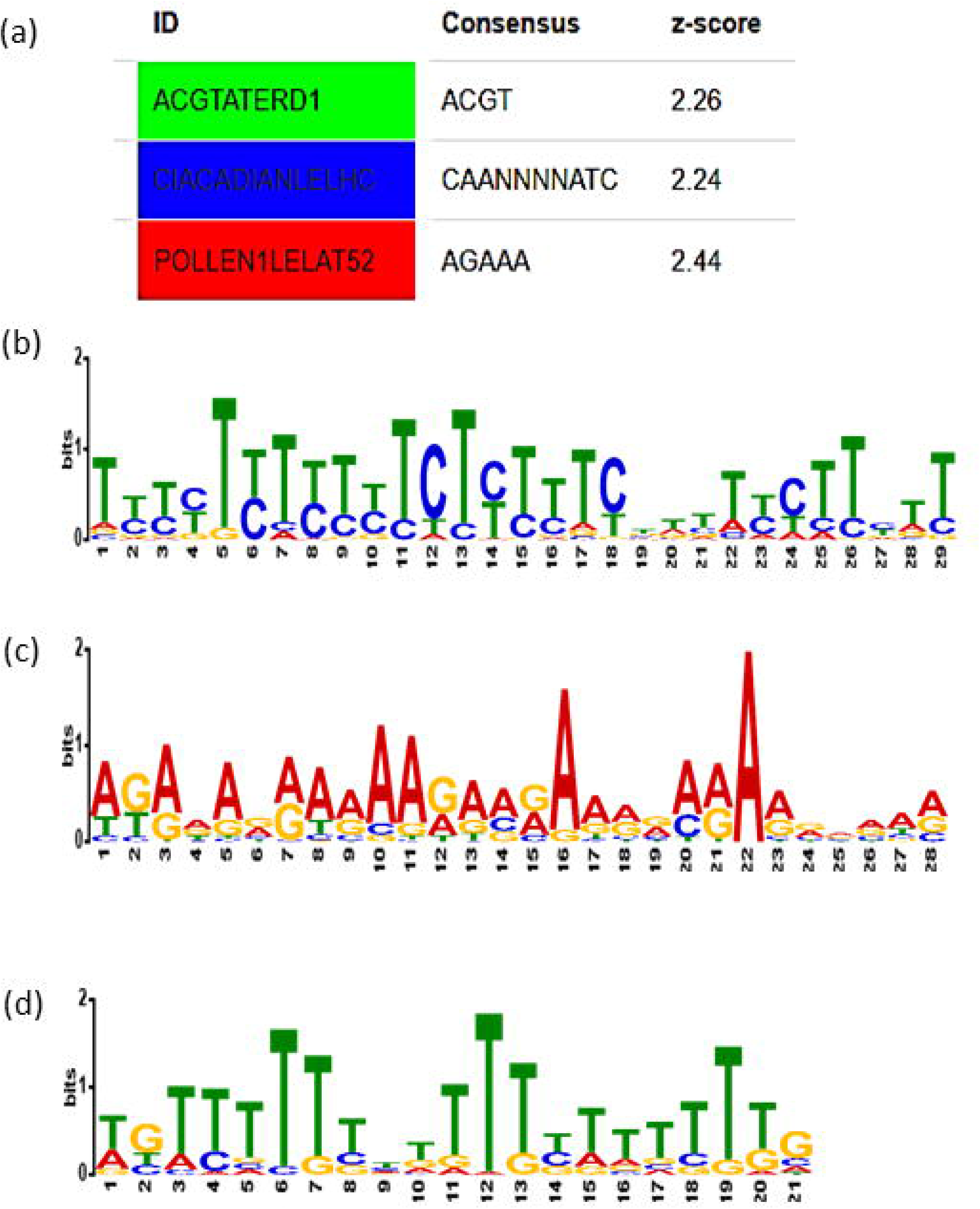
Conserved motifs in top 50 genes that co-expressed with *PIRL6* (a) Three known motifs and their consensus sequences overrepresented in the promoters based on Cistome analysis. (b) Conserved putative motif 1 generated by MEME Suite. (c), Conserved putative motif 2 generated by MEME Suite. (d), Conserved putative motif 3 generated by MEME Suite. X axis in the b, c, and d is the number of nucleotides indicating the length, Y axis indicating the bits (relative positional frequency of a particular nucleotide).

### Prediction of novel motifs in the promoters of *PIRL6* and its co-expressed genes

Three novel conserved motifs (M1, M2, and M3) were predicted from the promoters of 51 coexpressed genes (S. Fig. 6), by using the MEME web server. A total of 149 novel motifs (M1, M2, and M3), 29, 28, and 21 nucleotides-long (Fig. 5b), respectively, were detected in these promoters. While M1 was least common, present only in 46 of 51 (92%) genes analyzed, the other two were present over all the promoters analyzed. The negative strands showed a substantially larger distribution of motifs than the positive strands. Higher concentration of motifs (54 %) was found between −500 to −1 bp, upstream of the translation start sites.

In order to determine whether these putative motifs resembled any of the previously identified regulatory motifs for transcription factor binding, the *de novo* motifs were compared to registered motifs available in the pubic databases using the web application TOMTOM. CIS BP 2.0 single base pair DNA of *A. thaliana* was selected as the database to carry out the comparative study in the TOMTOM web application. M1, M2, and M3 matched with 86, 78, and 73, respectively, of the 872 known motifs that were available in the database. The top-matched motifs were chosen based on their estimated statistical significance levels. While the best possible candidate predicted for binding with both M1 and M2 belonged to the transcription factor C2C2dof_tnt.AT5G66940_col_a_m1 (AT5G66940) of the C2C2-DOF family, the best match for M3 was ABI3VP1_tnt.VRN1_col_a_m1 (VRN1) belonging to the ABI3-VP1 transcription factor family. The AT5G66940-encoded protein is a nuclear localized DNA-binding with one finger (DOF)-domain binding transcription factor. The DOFs are a group of plant-specific transcription factors mostly involved in the development and functioning of the vascular tissues, and some are mobile proteins that possibly control short- or long-distance signaling (Le Hir and Bellini 2013). The VRN1 transcription factor is previously known to be involved in the regulation of the vernalization (Levy et al. 2002) and flower development from meristematic tissue (Adam et al. 2007). In the case of the *PIRL6* promoter, the putative M3 motif relatively matching to VRN1 transcription factor was present only in the PA1 fragment and not the PA2 and PA3 fragments, indicating its possible role in directing pollen-specific expression.

### Functional abscisic acid- and gibberellin-responsive motifs identified in *PIRL6* promoter

The *PIRL6* promoter sequence showed the presence of two putative hormone-responsive elements, the ABRE (abscisic acid-responsive) and P-box (gibberellin-responsive) common for all the three lengths lies at -474 bp and -372 bp upstream of ATG start site, when analyzed using the PlantCare tool (Fig. 6g). To verify their function, tobacco leaf agroinfiltration assay was carried out with *Agrobacterium* strain GV3101 harboring either *PIRL6* PA1 or pCAMBIA1301 vectors. The binary vector pCAMBIA1301 and *PIRL6* PA1 are both in the same background, except that the *GUS* in pCAMBIA1301 is under the P35S constitutive promoter, and hence, it was used as a positive control. A set of leaves were subsequently subjected to either gibberellic acid or abscisic acid treatment, in parallel, so that if the hormone-responsive elements in the PA1 promoter responded, that would lead to alteration in GUS expression. Though GUS expression was seen in all leaves infected with either of the bacterial strains, infection with *Agrobacterium* harboring pCAMBIA1301 resulted in the highest expression, followed by infection with pAJ*PIRL*1, and then by those treated with either of the hormones (Fig. 6a-6e). The GUS activity of PA1 promoter was reduced to 1.34 or 1.35 folds (Fig. 6f), after the treatment with gibberellic acid or abscisic acid, respectively. Though these values in PA1 are not significantly different from the treated ones, it could be possible that the *cis*-elements in PA1 are mildly sensitive to gibberellic acid and abscisic acid.

**Fig. 6.**
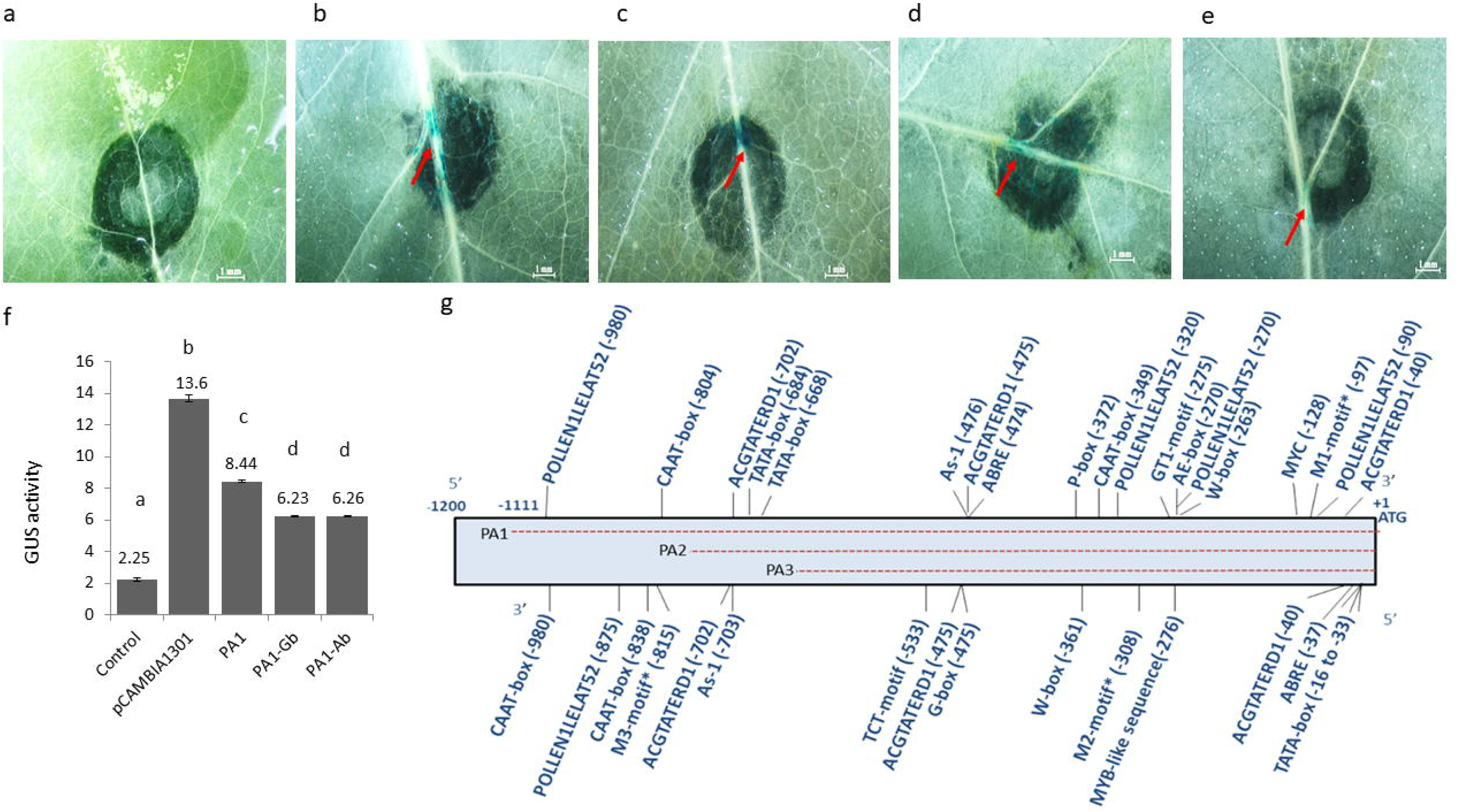
Transient expression of PA1::GUS in tobacco leaf and distribution of various *cis*-elements on *PIRL6* promoter. (a) Water treated tobacco leaf (control), (b) Tobacco leaf transformed with pCAMBIA 1301 , (c) Tobacco leaf transformed with PA1::GUS, (d) Absicissic acid-treated tobacco leaf transformed with PA1::GUS, (e) Giberillic acid-treated tobacco leaf transformed with PA1::GUS, (f) Quantitative measurement of GUS activity in control leaf (wild plant), leaf transiently transformed with pCAMBIA 1301, leaf transiently transformed with PA1, giberillic acid-treated tobacco leaf with PA1 promoter (PA1-Gb), absicissic acid-treated tobacco leaf with PA1 promoter (PA1-Ab), (g) distribution of various *cis*-elements such as P-box (pyrimidine box, gibberellin-responsive element) and ABRE (abscisic acid-responsive element) in the PA1 promoter of *PIRL6*, identified using PlantCARE. The red arrow shows the GUS positive blue tissue in the infected zone. The error bars indicate ± standard deviation of the means of four biological replicates. The symbols a, b, c and d denote a significant difference between the respective groups at p < 0.01, if not represented by the same symbol. GUS activity was assessed based on the nmol of 4-MU produced per minute per milligram of protein. Dotted line indicate the promoter fragments of different lengths. Asterisks indicate the predicted motif in promoter region whose function not confirmed.

## Discussion

Genetic engineering-related studies in plants have produced an appreciable amount of new information in the area of functional genomics, such as identifying the functions of various genes, promoters, and other regulatory elements, during the past two decades. Tissue-specific promoters are considered as a major discovery as they can be used as tools for targeted gene regulation, and thus have applications in crop improvement and metabolic engineering. Gametophyte-specific promoters are a type of tissue-specific promoters with wide application in plant breeding (Mithra et al. 2017), marker elimination from transgenic plants (Luo et al. 2007; Mlynarova et al. 2006; Verweire et al. 2007), and genome editing (Wang et al. 2015). Compared to the increasing range of crops undergoing these experiments, only a few gametophyte-specific promoters have been identified and confirmed to drive the expression of desirable genes. Information on *cis*-regulatory regions of gametophyte-specific promoters is also very less.

The model plant *A. thaliana* has always been a good resource of tissue-specific promoters. The male gametophyte-specific promoters from *A. thaliana* include those for the genes *AtSTP6* (Starke et al. 2003), *BCP1* (Xu et al. 1993), *AtVEX,* and *AtGEX2* (Engel et al. 2005). Similarly, some pollen-specific promoters reported from other plants include those driving the expression of genes *Zm908* from Maize, *LGC1* from lily, and *Os10g22450* from Rice (Smirnova and Kochetov 2020). Most of these promoters were discovered based on the reports of their tissue-specific expression. One important gene family to be involved in gametophyte development, extensively studied over the last decade in *Arabidopsis*, is *PIRLS*. This family comprises a total of nine genes. *PIRL6* is one of the main candidate from this family, initially reported to be involved in the development of pollen grains (McNichol 2016). Segregation distortion and a faulty pollen phenotype, which resulted in the RNAi knockdown experiment, strongly indicated the role of the *PIRL6* gene in pollen formation. Forsthoefel et al. (2018) reported that *PIRL6* is necessary for both male and female gametogenesis and proposed that unproductive alternative splicing adversely regulates the sporophytic expression of *PIRL6*. Hossain et al. (2022) conducted a promoter-*GUS* analysis of nine *PIRL* genes of *Arabidopsis*, including *PIRL6*. For all these genes, they analyzed the function of 2 kb-long fragment upstream of the ATG start codon. The 2 kb promoter of *PIRL6* controlled expression in both the male and female gametophytes. In this report we performed detailed deletion analysis of the *PIRL6* promoter, performed *in silico* comparisons of *PIRL6* with other co-expressed genes, and predicted some of the *cis*-acting elements.

Deletion analysis of promoters is the ideal method for a deeper understanding of promoter regions (Andersson and Sandelin 2020). Previously, such an approach has revealed the functions of the promoters of genes such as *AtUSP* (Bhuria et al. 2016), *PAL1* (Ohl et al. 1990), and *APETALA2* (Sharma et al. 2017) in *Arabidopsis*. We analyzed the expression of *GUS* reporter gene, regulated by three different lengths PA1, PA2, and PA3 (1128, 762, 646 bps) of *PIRL6* promoters. We identified the larger fragment, PA1 (-1111 to +17), to be essential for pollen-specific expression. Though we did not observe visible GUS expression in sporophyte and female gametophyte, GUS fluorimetric assay indicated a basal level expression in these tissues. We did not visualize GUS in female gametophyte probably because all our promoter versions were shorter than the one reported by Hossain et al. (2022).

Comparison of promoter sequences in co-expressing genes can predict conserved sequences and regulatory elements. Fifty genes that co-expressed with *PIRL6* were predicted to be invariably involved in cellular differentiation. Comparison of these 51 promoters showed the presence of three overrepresented elements POLLEN1LELAT52, ACGTATERD1 and CIACADIANLELHC. Of these, POLLEN1LELAT52 was a previously confirmed pollen-specific regulatory element of the tomato pollen-specific *lat52* gene (Filichkin et al. 2004). ACGTATERD1 motif is required for etiolation-induced expression of ERD1 (early responsive to dehydration) in *Arabidopsis* (Simpson et al. 2003). CIACADIANLELHC was reported as circadian rhythms-related transcription factor binding site in tomato (Piechulla et al. 1998). A previous study of *cis*-elements of various co-expressed genes in *Arabidopsis* male meiocytes also reported the over-representation of POLLEN1LELAT52 (Li et al. 2014). Apart from these, we predicted three novel motifs M1, M2, and M3 in *PIRL6* and its co-expressed genes. Of these, M3 was unique to PA1, as it was located at -815 bp upstream of the *PIRL6* translation start site. The intriguing part of this analysis is that this motif is predicted to be recognized mostly by VRN1 like transcription factor, which is generally involved in vernalization (Levy et al. 2002) and flower development from meristematic tissue (Adam et al. 2007). Though further confirmation of this prediction is required, there are no previous reports of VRN1 being involved in the regulation of genes in gametophytes.

Our tobacco leaf assay showed considerable transient expression of the PA1 promoter in sporophytic tissue, probably because the promoter-specificity was lost in the heterologous system. Nevertheless, this helped us in confirming the prediction about the presence of two hormone-responsive elements in the *PIRL6* promoter. The predicted ABRE ‘ACGTG’ (-474 bp) and Pbox element ‘CCTTTTG’ (-372 bp) were functional, as verified by the leaf assay. Treatment with either of these hormones downregulated PA1-GUS expression. Although both hormone-responsive elements are widely reported in promoters of different genes in different plants, ABRE comprises the most conserved sequence in dehydration-inducible promoters of genes such as At1g20440 (*ATCOR47*), Os11g0454300 (*RAB16A*), and Glyma04g01130 in *Arabidopsis*, rice, and soybean, respectively (Maruyama et al. 2012). Recently, Luo et al. 2022 reported the presence of ABRE element in the promoter of 69 out of 74 genes of the *TaGH9* family in wheat. Similar to our *PIRL6*, 69 of these genes expressed in male gametophyte and were involved in pollen fertility and anther dehiscence. Hence, it could be possible that ABA signaling is involved in pollen development and anther dehiscence, and this needs further validation.

Considering the increasing applications of gametophyte-specific promoters, especially in plant breeding, marker elimination and genome editing, it is important to verify their functional regions. Some of the predicted regulatory motifs can be used in the making of synthetic promoters. Thus the *PIRL6* promoter, which is essential for pollen-specific expression in *Arabidopsis* can also be used for such applications. While the requirement of applying these technologies to novel crops is increasing, it is also to be noted that promoters behave differently in different organisms. Hence, the *PIRL6* promoter can be tested in crops, where the previously reported promoters do not work. Novel promoters and *cis* regulatory elements can also be tested from the predicted genes which co-expressed with *PIRL6*.

## Conclusions

In this study we have characterized various lengths of the promoter region of *PIRL6* gene and identified a male gametophyte specific segment of length 1128 bp, which exclusively expressed in pollen grain. The GUS fluorimetric data confirmed the specificity of the promoter expression. Apart from that, we have predicted three novel potential *cis* acting elements over represented in the promoters of *PIRL6* co-expressing gene network. The functional validation of two hormone responsive element (ABRE and Pbox) of the *PIRL6* promoter also studied using tobacco leaf transient assay. To sum-up, the characterization of this tissue specific promoter fragment and *cis* acting element can be used in the field of transgenic technology, to develop synthetic promoter, and to study male sterility in plants.

## Acknowledgements

The authors would like to acknowledge Kerala State Council for Science, Technology and Environment (KSCSTE) for funding this research. Ajith acknowledges Central University of Kerala for his PhD fellowship.

## Statements and Declarations

### Author Contribution

JMS and ATG conceived the idea, designed the experiments and wrote the paper. ATG performed the experiments. All authors have read and approved the final manuscript

### Data availability

The datasets generated and analysed during this study are available from the authors on reasonable request.

### Competing Interests

The authors declare that they have no conflict of interest.

## Notes

### Competing Interest Statement

The authors have declared no competing interest.

