## Supplementary file 1 for "Functional profiling of the male gametophyte-specific promoter fragment of *Arabidopsis PIRL6* gene and prediction of *cis*-regulatory elements"

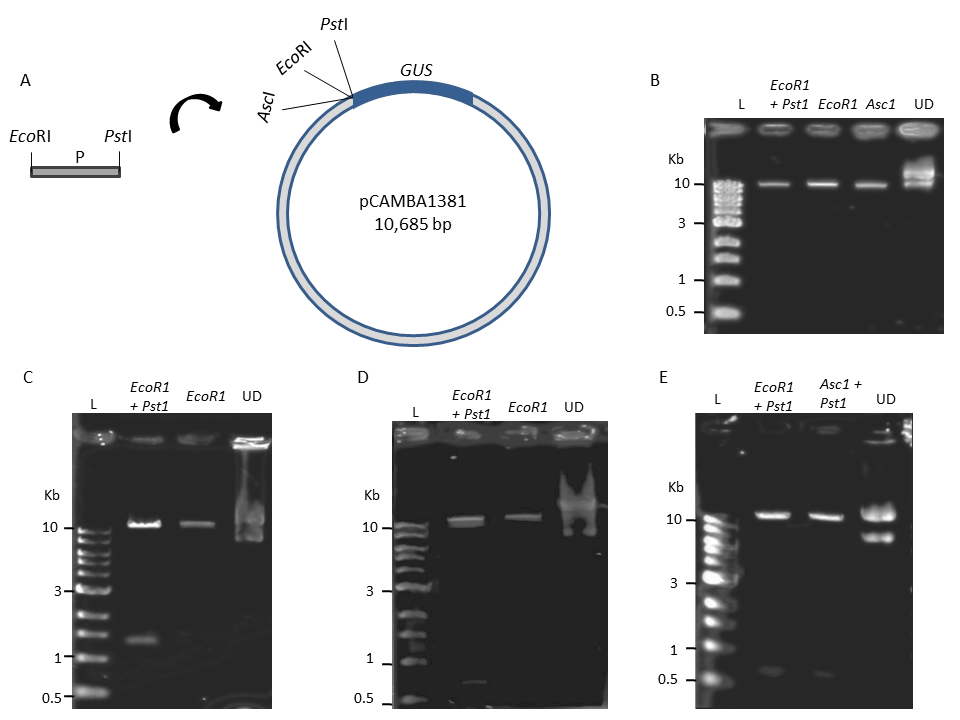

**S. Fig. 1** Cloning strategy and restriction digestion-based clone confirmation of the three promoters PA1, PA2 and PA3 into the multiple cloning site of pCAMBIA1381, upstream of the promoterless *GUS* gene. (A) Diagrammatic representation of the cloning of promoters denoted commonly as ‘P’. (A) Digestion of the vector pCAMBIA 1381 using various restriction enzymes (C) Confirmation of pAJPIRL1 with PA1. (D) Confirmation of pAJPIRL2 with PA2. (E) Confirmation of pAJPIRL3 with PA3. L is 1-kb ladder, UD is respective undigested vector/clone.

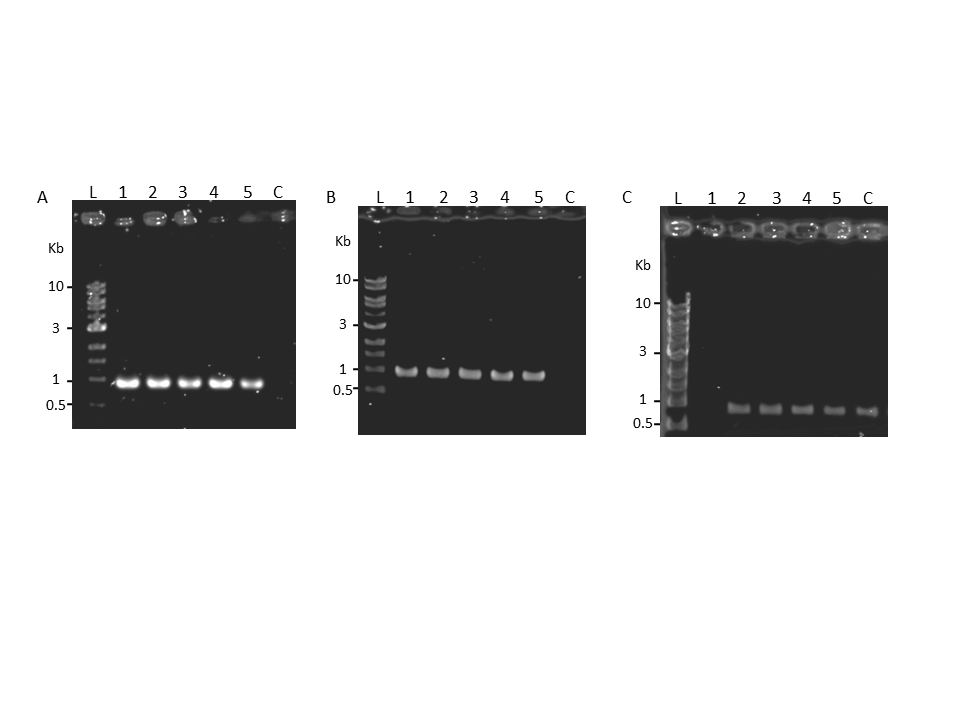

**S. Fig. 2** PCR confirmation of transformed plants using *GUS*-specific primers. (A) Plants transformed with pAJPIRL1. (B) Plants transformed with pAJPIRL2. (C) Plants transformed with pAJPIRL3. L is 1-kb ladder. The numbers 1 to 5 indicate the DNA of five independent transgenic plants from the respective transformations. C is the control DNA of untransformed plant.

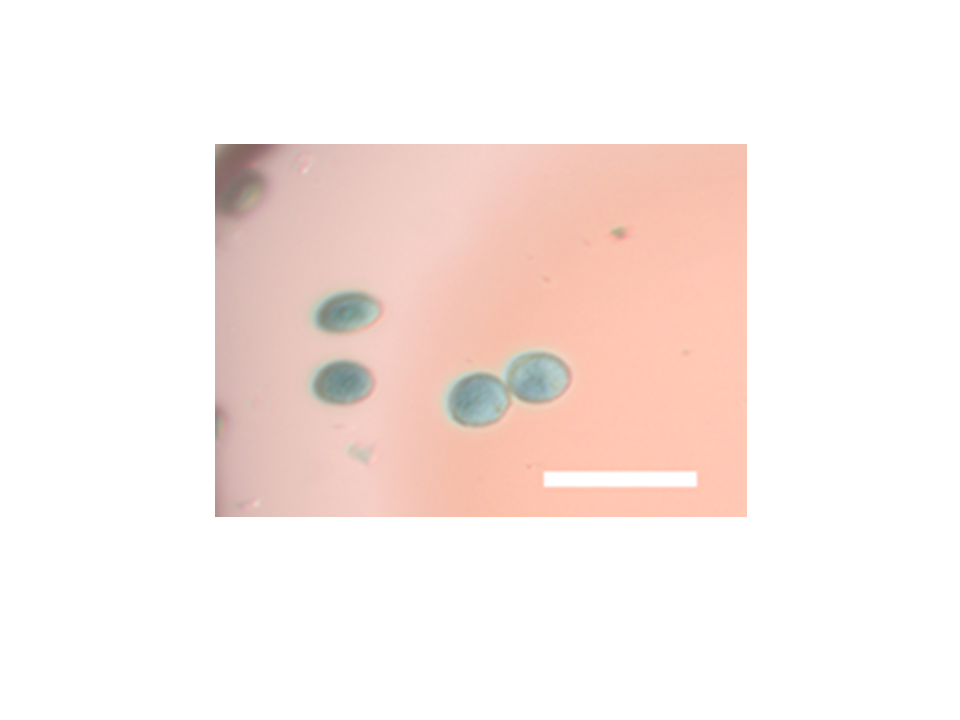

**S. Fig. 3** Pollen grains isolated after *GUS* staining from anther of PA1- *GUS* construct harboring *Arabidopsis* plant.

A
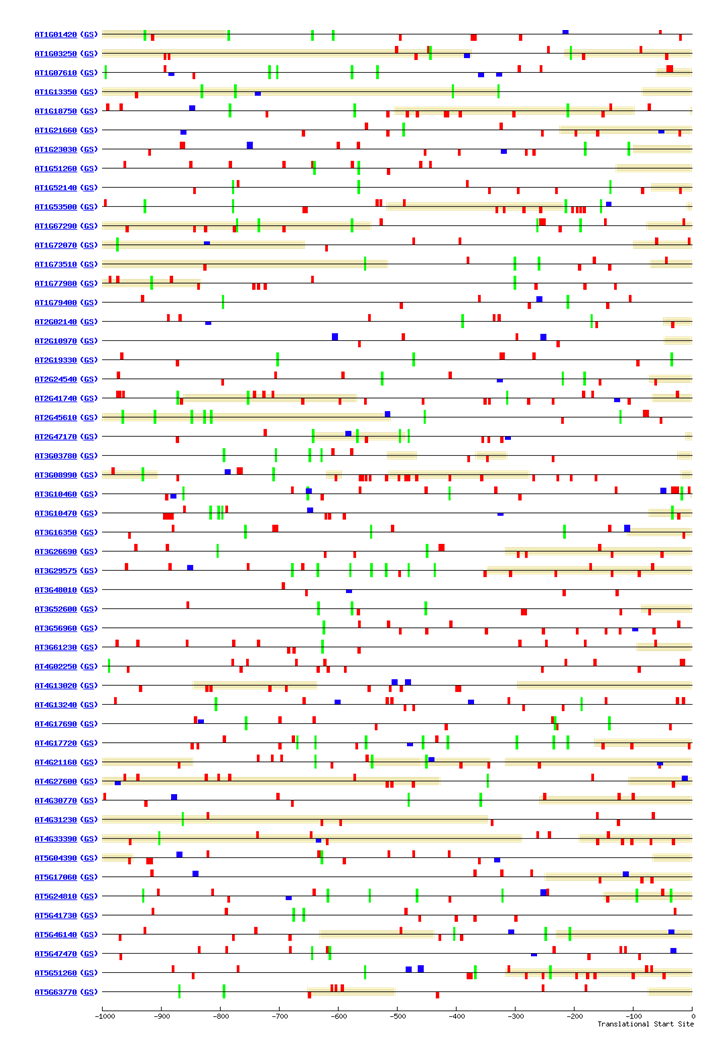

**S. Fig. 4** Cistome analysis of the promoters of genes that coexpressed with *PIRL6* (At2g19330). The red, blue and green boxes represent POLLEN1LELAT52, CIACADIANLELHC and ACGTATERD1 motifs, respectively. The numbers 0 to -1000 represent distance in base pairs from the translational start site. The presence of colored boxes on upper and lower side of the lines, represent their presence on positive and negative strands, respectively. The tan bars correspond to UTR sequences*.*

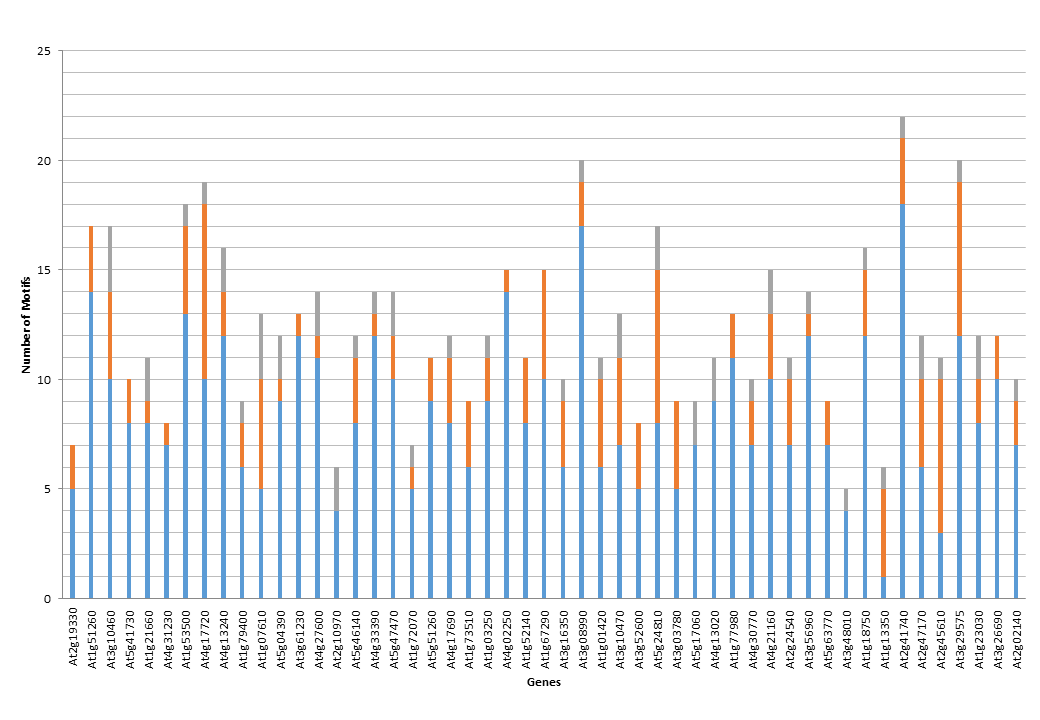

**S. Fig. 5** Distribution of the three overrepresented *cis* motifs in the gene network that co-expressed with *PIRL6* (At2g19330), generated during the Cistome analysis. The regions coloured in grey, red and blue indicate CIACADIANLELHC, ACGTATERD1, and POLLEN1LELAT52, respectively.

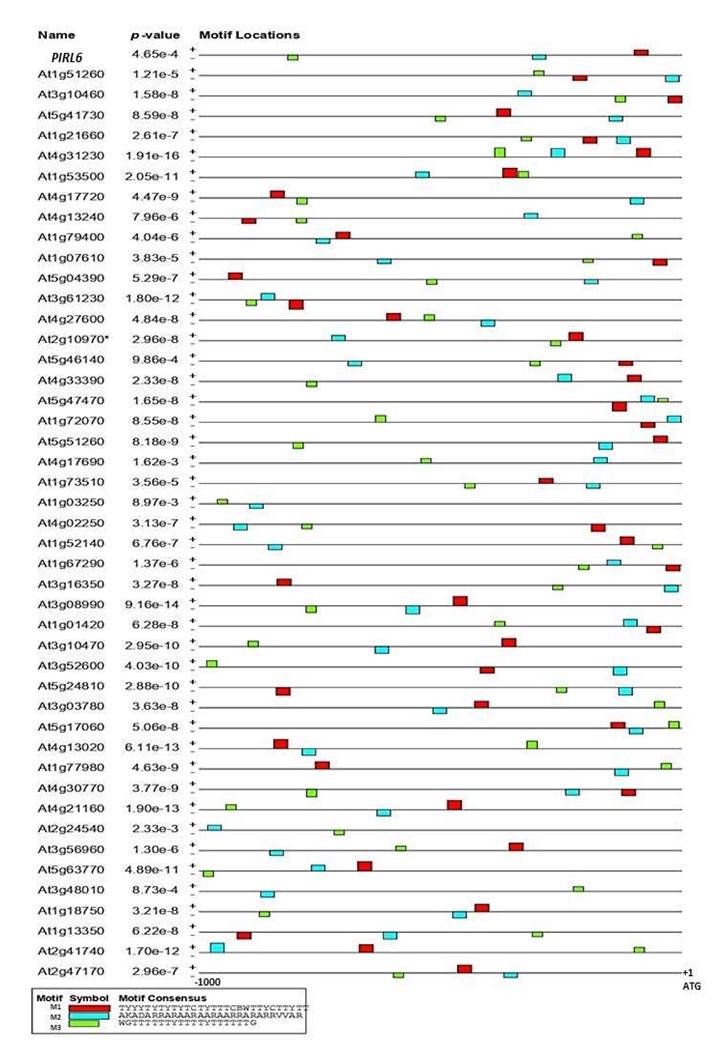

**S. Fig. 6** Distribution pattern of putative motifs predicted from the promoters (1 kb upstream of ATG start codon) of *PIRL6* (At2g19330) and its top 50 co-expressed genes, using MEME suite. The presence of colored boxes on upper and lower side of the lines, represent their presence on positive and negative strands, respectively.
