## Supplementary file 2 for "Functional profiling of the male gametophyte-specific promoter fragment of *Arabidopsis PIRL6* gene and prediction of *cis*-regulatory elements"

| Promoter/ gene name | Primer sequence (5’-3’) | | TM (°C) |
| --- | --- | --- | --- |
| PA1 | F | ACTTGAATTCTGGCCATGAATGGATGTTCT | 70 |
|  | R* | ACTTCTGCAGCTCCTCGCATATCATCTTT | 71 |
| PA2 | F | ACTTGAATTCCTCCACGAACAAAGCTAACAAAG | 70 |
|  | R* | ACTTCTGCAGCTCCTCGCATATCATCTTT | 71 |
| PA3 | F | ACTTGAATTCCTTGATCTCACTCACCTCGAAC | 71 |
|  | R* | ACTTCTGCAGCTCCTCGCATATCATCTTT | 71 |
| *GUS* | F | GGCAAAGTGTGGGTCAATAATC | 64 |
|  | R | CACGCGCTATCAGCTCTTTA | 65 |

* Common for PA1, PA2 and PA3

**S. Table 1** Details of primers and their annealing temperatures during PCR.

| No. | Accession no. | Gene name |
| --- | --- | --- |
| 1. | At2g19330 | *PIRL6* |
| 2. | At1g51260 | *LPAT3* |
| 3. | At5g41730 | *MLKL2* |
| 4. | At1g21660 | *AUXILIN-LIKE3* |
| 5. | At4g31230 | NA |
| 6. | At1g53500 | *ATMUM4* |
| 7. | At4g17720 | *BPL1* |
| 8. | At4g13240 | *ATROPGEF9* |
| 9. | At1g79400 | *ATCHX2* |
| 10. | At1g07610 | *MT1C* |
| 11. | At5g04390 | NA |
| 12. | At3g61230 | *ATPLIM2C* |
| 13. | At4g27600 | *NARA5* |
| 14. | At2g10970 | NA |
| 15. | [AT5G46140](https://www.arabidopsis.org/servlets/TairObject?id=132955&type=locus) | *ATDOA13* |
| 16. | At4g33390 | NA |
| 17. | At5g47470 | NA |
| 18. | At1g72070 | NA |
| 19. | At5g51260 | NA |
| 20. | At1g03250 | NA |
| 21. | At1g67290 | *GLOX1* |
| 22. | At3g16350 | *NID1* |
| 23. | At3g08990 | NA |
| 24. | At1g01420 | NA |
| 25. | At3g10470 | NA |
| 26. | At3g52600 | NA |
| 27. | At5g24810 | *ABC1K11* |
| 28. | At3g03780 | *ATMS2* |
| 29. | At5g17060 | NA |
| 30. | At4g13020 | NA |
| 31. | At1g77980 | *AGL66* |
| 32. | At4g30770 | NA |
| 33. | At4g21160 | *ZAC* |
| 34. | At2g24540 | *AFR* |
| 35. | At3g56960 | NA |
| 36. | At5g63770 | *ATDGK2* |
| 37. | At3g48010 | *ATCNGC16* |
| 38. | At1g18750 | NA |
| 39. | At1g13350 | *PRP4KB* |
| 40. | At2g41740 | NA |
| 41. | At2g47170 | NA |
| 42. | At2g45610 | NA |
| 43. | At3g29575 | *AFP3* |
| 44. | At1g23030 | *PUB11* |
| 45. | At3g26690 | *ATNUDT13* |
| 46. | At2g02140 | *LCR72* |
| 47. | At3g10460 | NA |
| 48. | At4g17690 | NA |
| 49. | At1g73510 | NA |
| 50. | At4g02250 | NA |
| 51. | At1g52140 | NA |

NA - Names of these genes are not available

**S. Table 2** Top 50 Tissue Specific co-expressed genes of *PIRL6* identified using the Expression Angler custom tool.
